# Dissociable episodic memory performance and subjective experience across the aphantasia imagery spectrum

**DOI:** 10.64898/2026.10.22.761078

**Authors:** William Duckett, Jon S. Simons

**Author notes:** Correspondence: Please address correspondence to Professor Jon Simons.

## Abstract

Mental imagery is often assumed to support episodic memory, yet whether this relationship depends on conscious access to imagery, and how reduced imagery differs from total aphantasia, remains unclear. The present study compared objective episodic memory performance, subjective trial-by-trial vividness, confidence, and response times in people reporting normal imagery, reduced imagery, or total aphantasia, across younger and older age groups. Participants completed an incidental-encoding episodic memory task requiring identification of visual or spatial changes in targets within scenes, alongside trial-by-trial vividness and confidence ratings. Across the full imagery spectrum, trial-level fluctuations in vividness were related to memory performance, whereas trait-level and averaged vividness measures were not. Moreover, reduced imagery and total aphantasia showed dissociable patterns of episodic memory performance and subjective experience. In the reduced-imagery group, vividness was a stronger predictor of accuracy and response times, vividness judgements were slower, and confidence was lower, suggesting greater metacognitive uncertainty when imagery is weak and inconsistent. Controls outperformed reduced and aphantasic groups on visual but not spatial changes, and both groups responded more slowly than controls; confidence did not differ between control and aphantasic groups. Age-related memory decline was observed in the control and reduced groups but not in the aphantasic group. Exit questionnaire data highlighted limited awareness of aphantasia among typical imagers and a persistent belief among aphantasic participants that their memory was inferior. Together, these findings show that reduced imagery should be distinguished from total aphantasia rather than treating both as a single population, and that trial-level fluctuations in vividness reveal relationships with episodic memory that trait-level and averaged measures do not capture.

## Introduction

Mental imagery, the ability to form perceptual-like representations in the absence of external stimuli (Pearson et al., 2015), is a familiar aspect of conscious experience for most people. Imagery and mental reconstruction are core components of episodic memory retrieval (Wittmann & Şatırer, 2022), with the ability to “picture” or “see” a past event being thought of as indicators of accuracy. However, the definition of *aphantasia* by Zeman et al. (2015) as a condition of “reduced or absent mental imagery”, reflected by at or near floor scores on the Vividness of Visual Imagery Questionnaire (VVIQ) (Marks, 1973) has challenged this assumption. Despite lacking the subjective experience of visual mental imagery, aphantasics, who represent approximately 2-6% of the population (Zeman, 2024), demonstrate intact everyday functioning (Zeman, 2024), with largely unimpaired memory ability when compared to those who report mental imagery (Azañón et al., 2025; Knight et al., 2026; Milton et al., 2021; Reeder et al., 2024; Siena & Simons, 2024). Consequently, the questions of how subjective mental imagery experience relates to objective memory performance, and how memory differs between those who do and do not experience mental imagery, require further investigation.

### Between-group differences

Across visual working memory and mental rotation tasks, individuals with aphantasia typically perform equivalently to those who report visual mental imagery (referred to as ‘controls’ hereafter) (Jacobs et al., 2018; Pounder et al., 2022; Reeder et al., 2024; Reeder & Figueroa, 2022; Zeman et al., 2015), with some also noting superior performance for aphantasics (L. Kay et al., 2024) and no discernible differences at the neurological level between groups (Knight et al., 2026). Notably, aphantasics also frequently demonstrate increased reaction times despite equivalent or greater accuracy (L. Kay et al., 2024; J. Liu & Bartolomeo, 2023; Pounder et al., 2022), which is often attributed to usage of compensatory non-visual strategies (Keogh et al., 2021). For controls, it has been reported that the time taken to rate vividness itself varies, with higher vividness ratings being given more quickly across 5 datasets, the effect of which increases with higher trait vividness (Barnett et al., 2026), as measured by self-report measures such as the VVIQ, suggesting response time may reflect the metacognitive process of rating one’s own vividness, alongside compensatory strategies. Furthermore, consistent patterns have been found during autobiographical recall, where aphantasics frequently report fewer and less rich details (Monzel et al., 2022, 2024; Pearson, 2019), which also extends to imaging future events (D’Argembeau & Van Der Linden, 2006; Dawes et al., 2022). Whether reduced details during autobiographical recall reflects an objective deficit in memory ability, or suggests those with aphantasia simply volunteer fewer details when asked, remains debated (Arcangeli, 2026; Dando et al., 2023; Michel et al., 2025), with some suggesting that mental imagery acts as an “emotional amplifier” (Abdelrahman et al., 2025) enhancing our ability to recall rich episodic details, without memory accuracy depending upon it.

However, when investigating the objective episodic memory performance of aphantasics, findings are less consistent. Several studies report no significant differences in long-term episodic memory performance between aphantasics and controls (Bainbridge et al., 2021; Milton et al., 2021; Palombo et al., 2015; Siena & Simons, 2024; Wittmann & Şatırer, 2022; Zeman et al., 2010), while others have found that aphantasics perform worse (Monzel et al., 2022) or better (Duckett & Simons, 2026) than controls. Dando et al. (2023) found that aphantasics recalled fewer correct details than controls during a mock eye-witness paradigm without making any more errors, suggesting a reduction in the quantity, but not quality, of memory for those with aphantasia. Additionally, Michel et al. (2025) and Phillips (2026) suggest that using forced-choice paradigms may be more appropriate when investigating the performance of aphantasics, with aphantasia perhaps being analogous to findings that those with blindsight may be able to process visual features despite no subjective insight into their performance (Michel et al., 2025). Taken together, objective memory performance appears largely preserved in those with aphantasia, albeit with increased response times and fewer autobiographical details. Given the inconsistency of the above findings, further direct investigation of episodic memory performance between those with aphantasia and controls with forced-choice paradigms is required. Therefore, the present study seeks to extend previous literature by using forced-choice methodology, with objective memory accuracy differences between those with aphantasia and controls as a primary outcome measure.

### Modality effects

Visual and spatial memory ability appear to be dissociable, with differences between controls and aphantasics being found. Bainbridge et al. (2021) found that when drawing previously studied images, participants with aphantasia recalled significantly fewer objects with less detail and colour compared to controls with typical imagery, but with equivalent spatial accuracy and fewer errors. This is consistent with evidence that visual-object and visual-spatial imagery may be separate (Blazhenkova, 2016; Blazhenkova & Kozhevnikov, 2010; Blazhenkova & Pechenkova, 2019), with spatial imagery being preserved in those with aphantasia despite lower visual imagery ratings (Keogh & Pearson, 2017), and spatial memory accuracy being equivalent (Crowder, 2018; Dawes et al., 2020; Keogh et al., 2021; Pounder et al., 2022; Reeder et al., 2024; Siena & Simons, 2024) or even superior (L. Kay et al., 2024) in comparison to controls. Moreover, trait measures of vividness have failed to predict spatial task performance, with no significant association between VVIQ scores and spatial memory accuracy being noted (Duckett & Simons, 2026), and other self-reported trait measures of mental imagery ability also failing to relate to performance on visuospatial tasks (Baade et al., 2025). However, the possible dissociation between visual and spatial memory has received little systematic investigation within the context of long-term episodic memory in aphantasia. The present study therefore investigates episodic memory performance in individuals with aphantasia across visual and spatial modalities, in comparison to controls.

### Subjective experience of aphantasia

Beyond objective performance differences, the subjective experience of living without mental imagery represents an important and frequently overlooked aspect of aphantasia research. Aphantasics often report lower confidence in their memory ability (Monzel, Dance, et al., 2023; Monzel et al., 2024; Monzel, Vetterlein, et al., 2023; Wittmann & Şatırer, 2022) and during perceptual tasks (J. Liu & Bartolomeo, 2023), despite intact objective performance, though some studies have failed to support a confidence gap (Reeder et al., 2024). Moreover, for many aphantasics, discovering that they may have aphantasia is described as a critical “light-bulb” moment for one’s sense of self (Digard et al., 2026) and a significant aspect of one’s identity (Mawtus et al., 2024). Attitudes of aphantasics towards a lack of mental imagery vary considerably, from acceptance to self-doubt, and even attempts to find a “cure” for their experience (Digard et al., 2026), highlighting a heterogeneity which objective group comparisons fail to capture. The present study therefore also seeks to understand the subjective experience of individuals with aphantasia through free text and multiple-choice questions.

### Reduced access to mental imagery

Aphantasia itself may also be heterogeneous. The “reduced or absent” characterisation of aphantasia (Zeman et al., 2015) includes two distinct types of subjective experience: a total lack of mental imagery (associated with the lowest possible score of 16 on the VVIQ), and very weak but present access to imagery (VVIQ = 17-32). Sometimes referred to as “hypophantasia” (Reeder et al., 2024), it has been estimated that 24%-36% of aphantasics experience some visual mental imagery (Dawes et al., 2024), with the estimated prevalence for individuals with VVIQ scores of 17-32 being 3.1% (Dance et al., 2022). Cutoff scores for aphantasia vary in the literature, ranging from ≤23 (Zeman et al., 2020), to ≤32 (Cabbai et al., 2023; Dance et al., 2022; Siena & Simons, 2024; Wicken et al., 2021), resulting in many individuals with reduced imagery being collapsed into aphantasic samples, potentially obscuring effects. Indeed, Pounder et al. (2022) demonstrated a significant response time difference between those scoring exactly 16 on the VVIQ and controls, which was missed by categorising all ≤24 VVIQ scores as aphantasic. People with reduced imagery have since been shown to differ from both aphantasics and controls in reasoning (Le Clézio et al., 2026), implicit priming (Purkart et al., 2025), and even mental health outcomes (Kvamme et al., 2026). Whether differences between those with reduced imagery, aphantasics, and controls extend to episodic memory performance remains largely unexplored. Analysis of data from Siena and Simons (2024) by Duckett and Simons (2026) found no significant difference in episodic memory performance between those with reduced imagery and those with aphantasia. However, this comparison was not the primary focus of either study, and larger sample sizes are needed to disentangle the two groups, therefore the present study was designed to consider those with reduced visual mental imagery, alongside those with aphantasia and controls.

### State, not trait vividness

Another question which requires further investigation is to what extent vividness, as a continuous measure, predicts memory performance in those with typical imagery. Several studies report no relationship between memory performance and both trait VVIQ scores (Fernandes et al., 2018; Marre et al., 2021; Thorudottir & Sigurdardottir, 2021) and averaged trial-by-trial vividness ratings (Duckett & Simons, 2026; Siena & Simons, 2024). However, these findings may reflect how vividness is measured and analysed, rather than whether a relationship is present. Trial-by-trial vividness ratings have been shown to predict subsequent memory performance where trait measures do not (D’Angiulli et al., 2013). More broadly, trial-by-trial state measures of vividness demonstrate higher predictive value and larger effect sizes than the VVIQ across behavioural, cognitive, and neurological studies (Runge et al., 2015, 2017), an effect which has also been observed for imagery-based task performance (Z. Liu et al., 2025), and metacognitive discrimination between imagery and perception (Dijkstra & Fleming, 2023). Notably, while Duckett and Simons (2026) found no relationships between averaged trial-by-trial vividness ratings or VVIQ scores and episodic memory performance in controls, within-subject trial-by-trial vividness ratings were associated with probability of correct memory responses. The present study therefore seeks to implement trial-by-trial ratings of vividness to capture state-level fluctuations of vividness, and to examine if this relationship extends to reduced and absent imagery groups.

### Age-related effects

While trial-by-trial state measures of vividness provide insight into the relationship between mental imagery and memory, the nature of this relationship may vary across the lifespan. Memory decline in healthy aging has been well-documented (De Luca et al., 2003; Hedden & Gabrieli, 2004). If visual memory is dependent, at least in part, on visual mental imagery (Keogh et al., 2021; Palombo et al., 2015), it might be expected that age-related memory decline may be accompanied by reduced imagery vividness. However, a dissociation in older adults has been noted, with subjective vividness ratings being higher despite objectively worse memory performance (Cooper & Ritchey, 2022; Duckett & Simons, 2026; Faul et al., 2025; Folville et al., 2020, 2021), alongside a greater variability in vividness ratings within older adults (Folville et al., 2021). Not all evidence supports this dissociation, however, as Mojescik et al. (2024) found that both younger and older adults anchor their subjective vividness ratings to conceptual gist rather than perceptual detail, with lower vividness ratings in older adults mirroring lower memory performance. Additionally, Gulyás (2022) found that average VVIQ scores decline from ages 14 to 55, not unlike the age-related decline seen in memory performance. Taken together, current findings demonstrate that the conditions under which vividness and memory begin to dissociate across the lifespan are unclear. Moreover, this relationship has not been examined in individuals with aphantasia or reduced imagery. The present study therefore compares younger and older adults with aphantasia, reduced imagery, and controls to examine age-related differences in subjective vividness and objective memory performance.

### Hypotheses

Regarding the relationship between vividness and memory accuracy, it is hypothesised that within-subject trial-by-trial vividness ratings will significantly predict memory accuracy (*H1)*, with trait and averaged measures demonstrating weaker relationships with memory accuracy than state measures (*H2*). Concerning group differences, those with aphantasia and reduced imagery are hypothesised to demonstrate superior accuracy in spatial memory modalities than controls (*H3*), with lower accuracy in visual modalities (*H4*), alongside longer response times (*H5*), and lower confidence ratings for participants with aphantasia specifically (*H6*). Higher vividness ratings are hypothesised to be associated with shorter response times when making vividness judgments (*H7*). Finally, regarding age-related effects, younger participants are hypothesised to outperform older participants in memory accuracy across imagery groups (*H8*), with older adults hypothesised to demonstrate a weaker relationship between vividness and memory performance (*H9*).

## Methods

### Participants

A total of 120 participants (59 Male, 61 Female) were recruited from the United Kingdom and United States of America, with English being a first or native language. 60 participants (30 Male, 30 Female) with typical mental imagery ability (VVIQ 33-80) were recruited via the online research platform *Prolific*. An additional 3 participants were excluded due to failing attentional checks. 60 participants (29 Male, 31 Female) with self-reported aphantasia were recruited via the *Aphantasia Network*. An additional 6 participants were recruited from the *Aphantasia Network* but were excluded due to either being outside the UK or US, and/or not having English as a first or native language. Written informed consent was obtained in a manner approved by the University of Cambridge Psychology Research Ethics Committee, and the research was conducted in accordance with the Declaration of Helsinki.

Upon inspection of the data, 4 participants with self-reported aphantasia had VVIQ scores exceeding 32: a commonly accepted cutoff for aphantasia. As a result, these participants were excluded from analyses, resulting in 116 total participants. To investigate effects of reduced, but not totally absent, mental imagery ability, those who reported VVIQ scores of >16 but ≤32 were categorised into the “Reduced” mental imagery ability category. This resulted in 3 mental imagery ability groups: Control (VVIQ >32, N= 60), Reduced (VVIQ 17-32, N = 20), and Aphantasic (VVIQ = 16, N = 36). A histogram of VVIQ score distributions can be seen in Figure 1.

**Figure 1.**
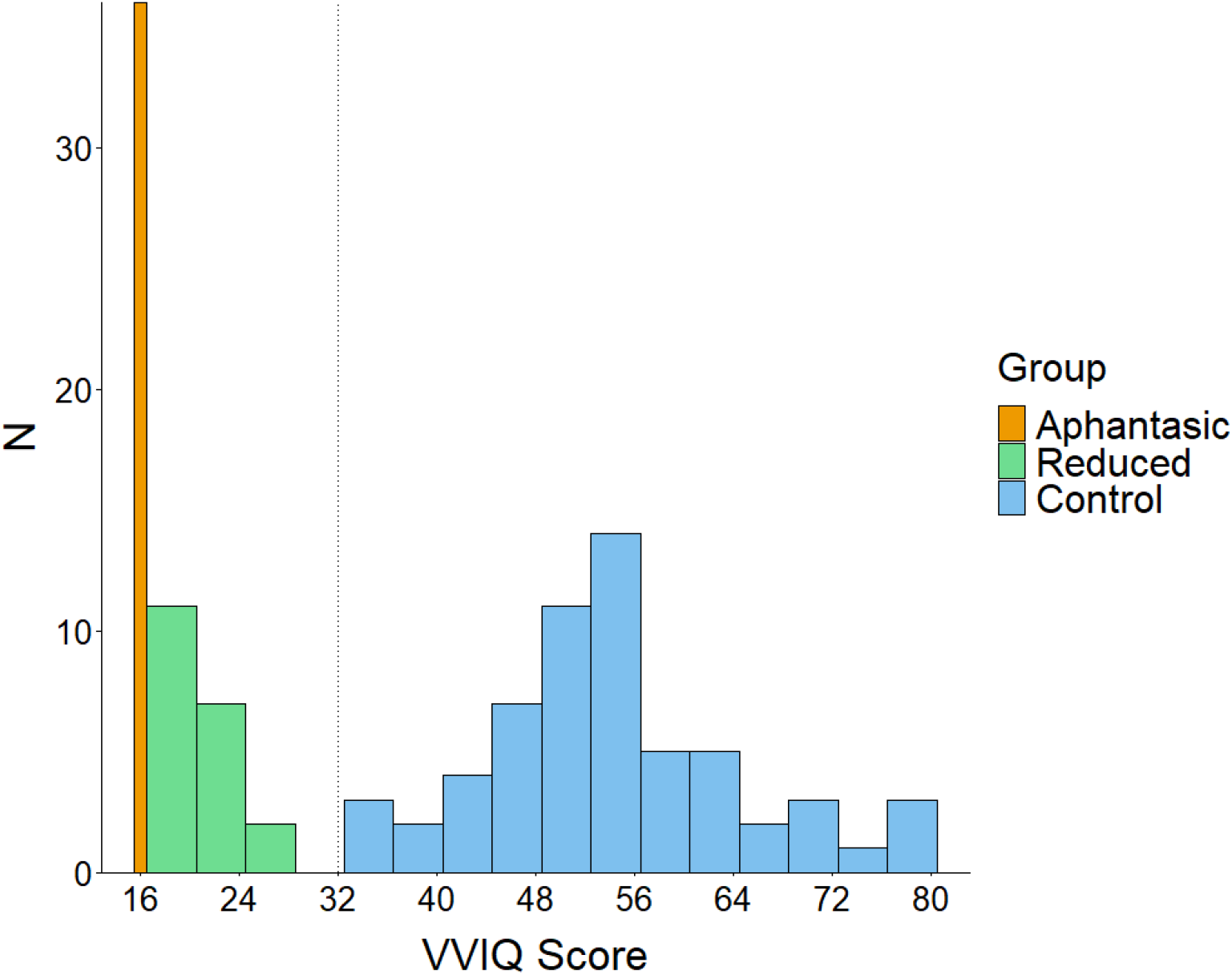
Histogram of VVIQ scores. Dashed vertical line indicates Control imagery cutoff score of 32.

To investigate potential age-related effects, recruitment aimed to obtain participants aged 18-30 (Younger) and 60-75 (Older). For the Control imagery group, 30 participants were aged 18-30 (Mean 25.6, 15 Female, 15 Male) and 30 aged 60-72 (Mean = 64.9, 15 Female, 15 Male). For the Aphantasic group, 16 participants were aged 19-30 (Mean = 25.4, 8 Male, 8 Female) and 20 were aged 60-75 (Mean = 66.7, 13 Female, 7 Male). For the Reduced imagery group, 12 participants were aged 18-29 (Mean = 23.4, 4 Female, 8 Male) and 8 were aged 61-74 (Mean = 67.0, 1 Female, 7 Male).

### Materials

Stimuli were adapted from Duckett and Simons (2026), consisting of 40 images of scenes, such as rooms, parks, and gardens each containing a target object within them. Two duplicate versions of each scene were created, one with a red circle to highlight the target object, and another with a visual (colour) change, spatial (location) change, or no change applied to the target object. Greyscale images of each target object were also created. Each scene and target were used only once. In total, there were 14 visual change, 13 spatial change, and 13 no change stimuli, see Figure 2 for example stimuli.

**Figure 2.**
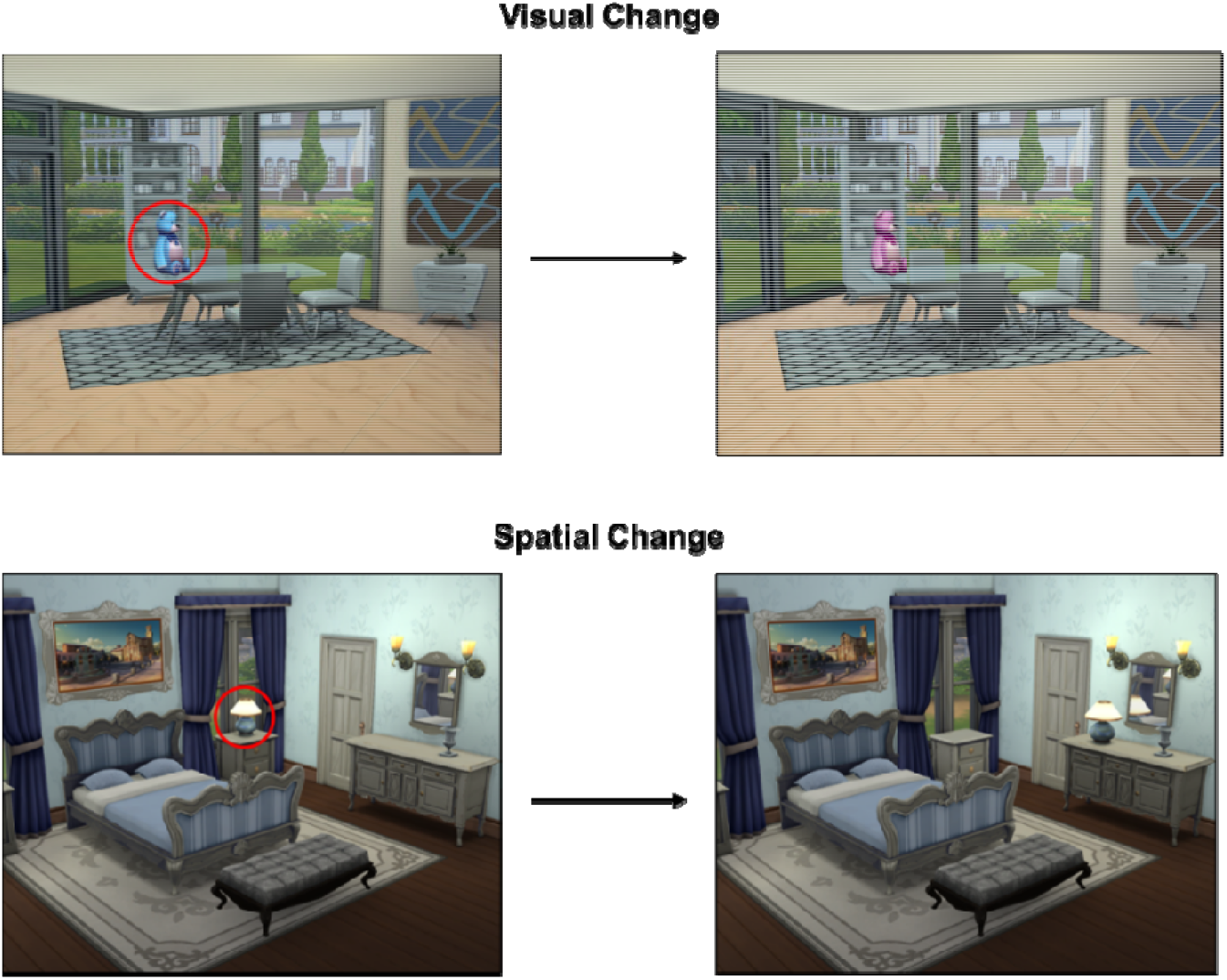
Example stimuli acquired from Duckett and Simons (2026).

Participants were also required to complete the VVIQ (Marks, 1973) to assess trait-level mental imagery vividness. Instruction graphics used to explain the concept of mental imagery were acquired and used with permission from the *Aphantasia Network*.

Upon completion of the study, depending on imagery group (Control, or Reduced and Aphantasic), participants were asked to complete a set of optional questions related to mental imagery, see Appendix A for a full list of questions.

### Design

The study followed a quantitative design. Dependent variables included memory accuracy, VVIQ score, trial-by-trial vividness ratings, trial-by-trial confidence ratings, and trial-by-trial response times. Independent variables included imagery group (Aphantasic, Reduced, or Control), age category (Younger or Older), and change modality (Visual, Spatial, or None). Measurements allowed for both between group and repeated measures analyses.

Although free-text responses were collected, the number of respondents was insufficient for formal qualitative analyses.

### Procedure

The study followed an identical procedure to Duckett and Simons (2026), consisting of two phases. The first Encoding phase followed an incidental encoding approach to ensure participants relied primarily on visual processing for subsequent memory judgements, avoiding strategies such as rehearsal and verbal scaffolding. Participants viewed stimuli for 5 seconds, followed by the same stimuli with the target object circled for a further 5 seconds. Participants were then asked two pseudorandomised questions related to visual and spatial aspects of the prior scene to act as a means for incidental encoding, with a fixation cross between each stimulus. The fixation cross would turn red pseudorandomly 20% of the time requiring the participants to press the Space Bar to continue, to act as an attentional check. The Encoding phase was split into 4 blocks of 10 stimuli in a within-block randomised order, resulting in 40 total trials. See Figure 3 (top) for a schematic representation of the Encoding procedure.

**Figure 3.**
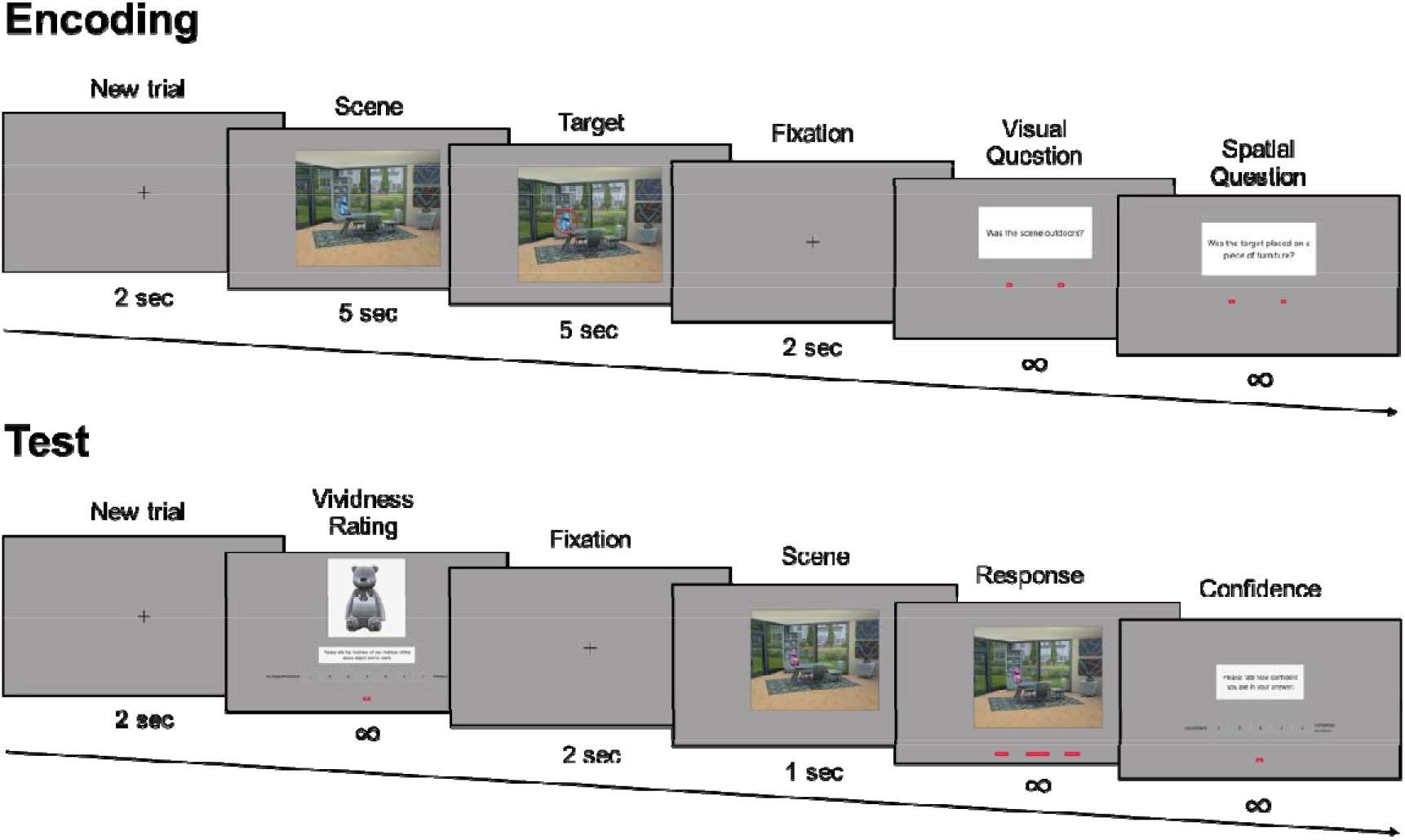
Diagram of Encoding (top) and Test (bottom) procedures, acquired from Duckett and Simons (2026).

Following the Encoding phase, participants were required to complete the VVIQ (Marks, 1973) before proceeding to the Test phase. In the Test phase participants were instructed that their memory of the previously presented scenes would be tested. Additionally, they were informed a change may have been applied to the target object: either a Visual (colour), Spatial (location), or No Change, and that it was their task to select from each respective option which they believe had occurred. Before making their decision, participants were shown a greyscale image of the target object and asked to rate their vividness (Ranging from 1: *No image whatsoever*, to 7: *Perfectly vivid*) of the memory of that scene and object and asked to rate their confidence (Ranging from 1: *Unconfident*, to 5: *Completely confident*) in their answer after making their decision. Images presented in block 1 of the Encoding phase would appear in block 1 of the Test phase, block 2 of Encoding in block 2 of Test, and so forth. See Figure 3 (bottom) for a schematic representation of the Test procedure.

## Results

### Data preparation

Data outputs were cleaned and analysed using R Statistical Software (v 4.4.2; R Core Team, 2024) with the *tidyverse* package (v 2.0.0; Wickham et al., 2019).

Test phase trials in which participants responded above the 95^th^ percentile of reaction times were excluded from analyses to account for extremely slow responses. This resulted in 232 total trials (8.0% of Aphantasic, 8.4% of Reduced and 2.1% of Control trials) being excluded from analyses, with 4408 total trials remaining. No participants failed >2 fixation cross attentional checks.

### Memory Performance

To investigate overall relationships between memory performance and vividness measures (VVIQ and mean trial-by-trial vividness ratings), a series of Spearman’s correlations were carried out as all measures failed Shapiro-Wilks tests for normality. Analyses revealed no significant correlation between memory performance and VVIQ scores (r(114) = 0.03, *p* = .752), nor between memory performance and mean trial-by-trial vividness ratings (r(114) = 0.06, *p* = .510). There was, however, a significant positive correlation (r(114) = 0.87, *p* = <.001) between mean trial-by-trial vividness scores and VVIQ. See Figure 4 for scatterplots of these relationships.

**Figure 4.**
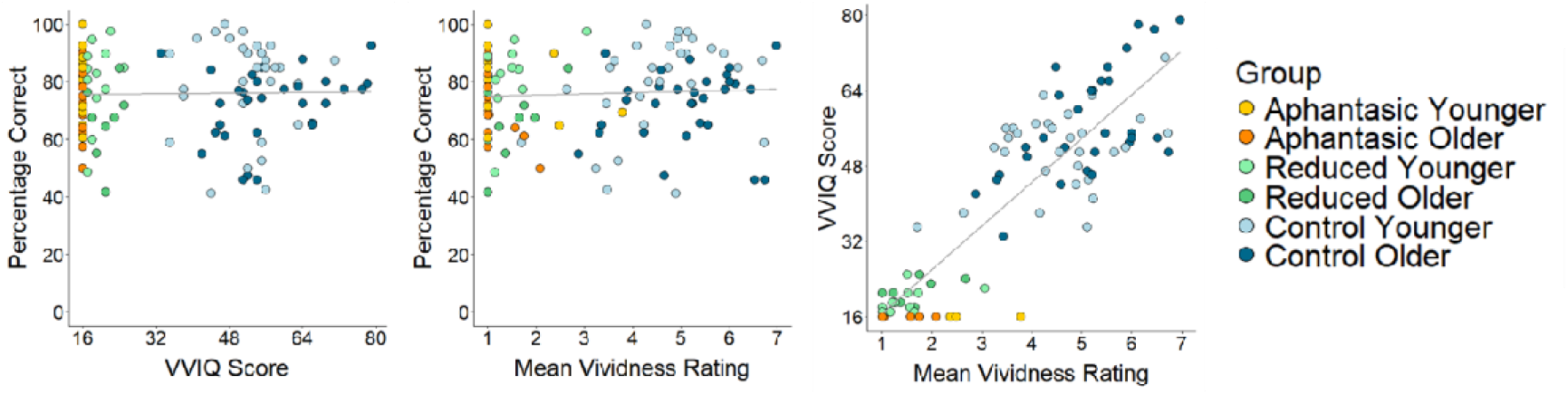
Scatterplots of the relationships between memory performance and vividness ratings (mean trial-by-trial vividness ratings and VVIQ score).

A two-way ANOVA was conducted to investigate the effects of imagery group (Aphantasic, Reduced, and Control), and age category (Younger or Older) on memory performance. Analyses revealed no significant effect of imagery group (*F*(2, 110) = 0.09, *p* = .912), but a significant effect of age group (*F*(1, 110) = 9.68, *p* = .002), on memory performance, suggesting similar performance across imagery groups, with Younger participants outperforming Older participants across groups. There was no significant interaction between imagery and age groups (*F*(2, 110) = 0.59, *p* = .559), indicating the effect of age was similar across imagery groups. Exploratory pairwise comparisons of estimated marginal means were conducted to compare performance of Younger and Older participants in each group. Results indicated Younger participants outperformed Older individuals, with a non-significant difference between age groups in the Aphantasic group (*p* = .180), but a significant difference in the Reduced (*p* = .026) and Control groups (*p* = .049). See Figure 5 for visualisations of memory performance by imagery and age group.

**Figure 5.**
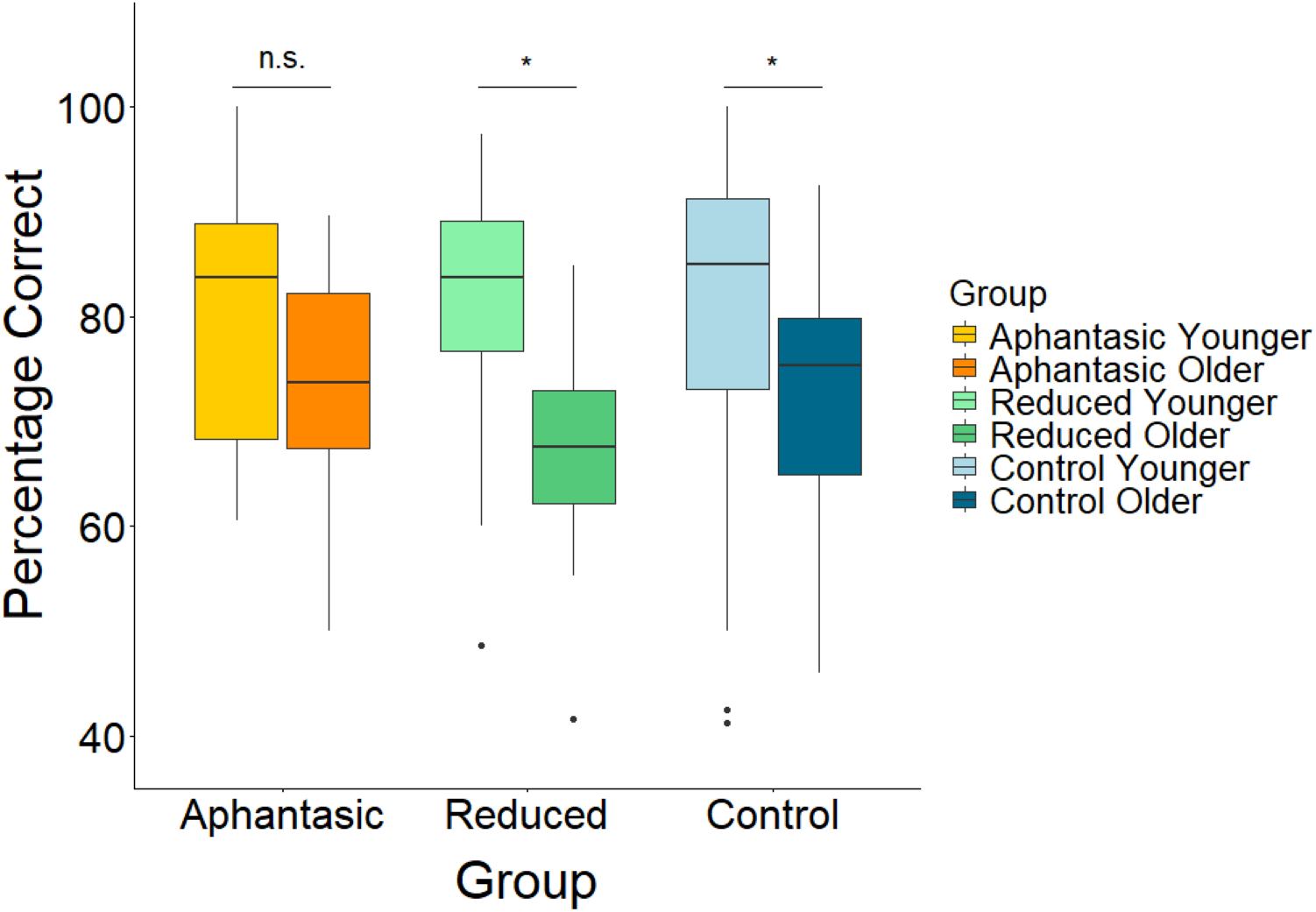
Boxplots of memory performance by imagery and age group. Significance lines denote pairwise comparisons of estimated marginal means.

While the data passed a Levene’s test for homogeneity of variance (F(5, 110) = 0.44, *p* = .822) a Shapiro-Wilks test revealed a non-normal distribution of residuals (W = 0.94, *p* = <.001). Therefore, a non-parametric aligned rank transform (ART) ANOVA was conducted using the *ARTool* (v 0.11.2; M. Kay et al., 2025) package in R. ART analyses revealed the same results with a non-significant effect of imagery group (*F*(2, 110) = 0.16, *p* = .851), a significant effect of age group (*F*(1, 110) = 10.56, *p* = .002), and a non-significant interaction between groups (*F*(2, 110) = 0.58, *p* = .561).

Analyses of memory performance up to this point used averaged, aggregated, memory scores and vividness ratings across 40 trials. In order to conduct analyses on a trial-by-trial basis, accounting for individual variability, a Binomial Generalised Linear Mixed Effects (GLMM) model using the *lme4* package (v 1.1.38; Bates et al., 2015) was conducted in R. To appropriately capture trial-by-trial variations of vividness for each participant, centring-within-cluster vividness scores were calculated for each participant, producing a trial-specific *Relative Vividness* measure. This was calculated by individually subtracting each participant’s average vividness rating across trials from vividness ratings given on each trial. To investigate the contribution of this trial-by-trial relative vividness measure, between-participant vividness measures (average trial-by-trial vividness and VVIQ), change modality, and imagery group on memory accuracy, the model was run with the following formula:

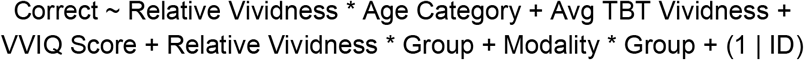

Where participant ID is entered as a random effect, with all others being entered as fixed effects. Interaction terms allow for interpretation of different groups’ use of vividness when making memory judgements, and performance differences between imagery groups in each modality. See Table 1 for statistical outputs.

**Table 1.** Statistical outputs following a GLMM predicting correct responses from other measures.

| Variable | Estimate | SE | p |
| --- | --- | --- | --- |
| (Intercept) | 1.28 | 0.6 | .034* |
| Relative Vividness | 0.26 | 0.05 | <.001*** |
| Age Category (Older) | -0.61 | 0.16 | <.001*** |
| Average Trial-by-Trial Vividness | 0.17 | 0.1 | .084 |
| VVIQ Score | 0.001 | 0.01 | .913 |
| Group (Aphantasic) | 0.86 | 0.49 | .076 |
| Group (Reduced) | 0.83 | 0.48 | .088 |
| Modality (Spatial) | -0.67 | 0.14 | <.001*** |
| Modality (Visual) | -0.8 | 0.13 | <.001*** |
| Relative Vividness * Age Category (Older) | 0.07 | 0.07 | .293 |
| Relative Vividness * Group (Aphantasic) | -0.07 | 0.15 | .636 |
| Relative Vividness * Group (Reduced) | 0.48 | 0.19 | .011* |
| Group (Aphantasic) * Modality (Spatial) | 0.13 | 0.23 | .575 |
| Group (Reduced) * Modality (Spatial) | -0.18 | 0.29 | .525 |
| Group (Aphantasic) * Modality (Visual) | -0.58 | 0.22 | .008** |
| Group (Reduced) * Modality (Visual) | -0.67 | 0.28 | .017* |

Note: Reference groups are Younger (Age Category), Control (Group), and No Change (Modality).

The above model initially produced convergence warnings; however, parameter estimates were highly consistent across optimisers when tested using allfit() from the *lme4* package (v1.1.38; Bates et al., 2015), indicating that the results were robust.

Results indicate that for Younger Control imagery participants, higher Relative trial-by-trial vividness significantly predicts better memory accuracy. Furthermore, a significant effect of age was found, with Younger participants performing significantly better than Older participants. The interaction between Relative Vividness and Age Category was not significant, indicating that the utility of vividness is similar across Younger and Older participants. A significant interaction between Relative Vividness and the Reduced imagery group was found, indicating that the effect of vividness in the Reduced group was stronger than in the Control reference group, with no significant difference in the effect of vividness between those in the Aphantasic group and the Control group. Moreover, a non-significant interaction for the Spatial modality was found for both Reduced and Aphantasic groups, but a significant negative interaction in the visual modality. This indicates similar performance across groups for No Change and Spatial stimuli, but significantly worse performance on Visual stimuli for those in the Reduced and Aphantasic group when compared to Control imagery ability participants.

To further investigate relationships between Relative trial-by-trial vividness ratings and accuracy, predicted probabilities were calculated using the above model. Analyses were conducted using the *ggeffects* (v2.3.1; Lüdecke, 2018) package in R to predict the probability of a correct response given a Relative Vividness rating for each age category and imagery group. As shown in Figure 6, higher Relative Vividness ratings were associated with greater accuracy, with the clearest difference evident across imagery groups. However, the extrapolated predictions for the Aphantasic and Reduced imagery groups should be interpreted with caution due to a lack of observed data in extreme Relative Vividness ratings in these groups. See Appendix B for a table of frequencies of each Relative Vividness rating by group.

**Figure 6.**
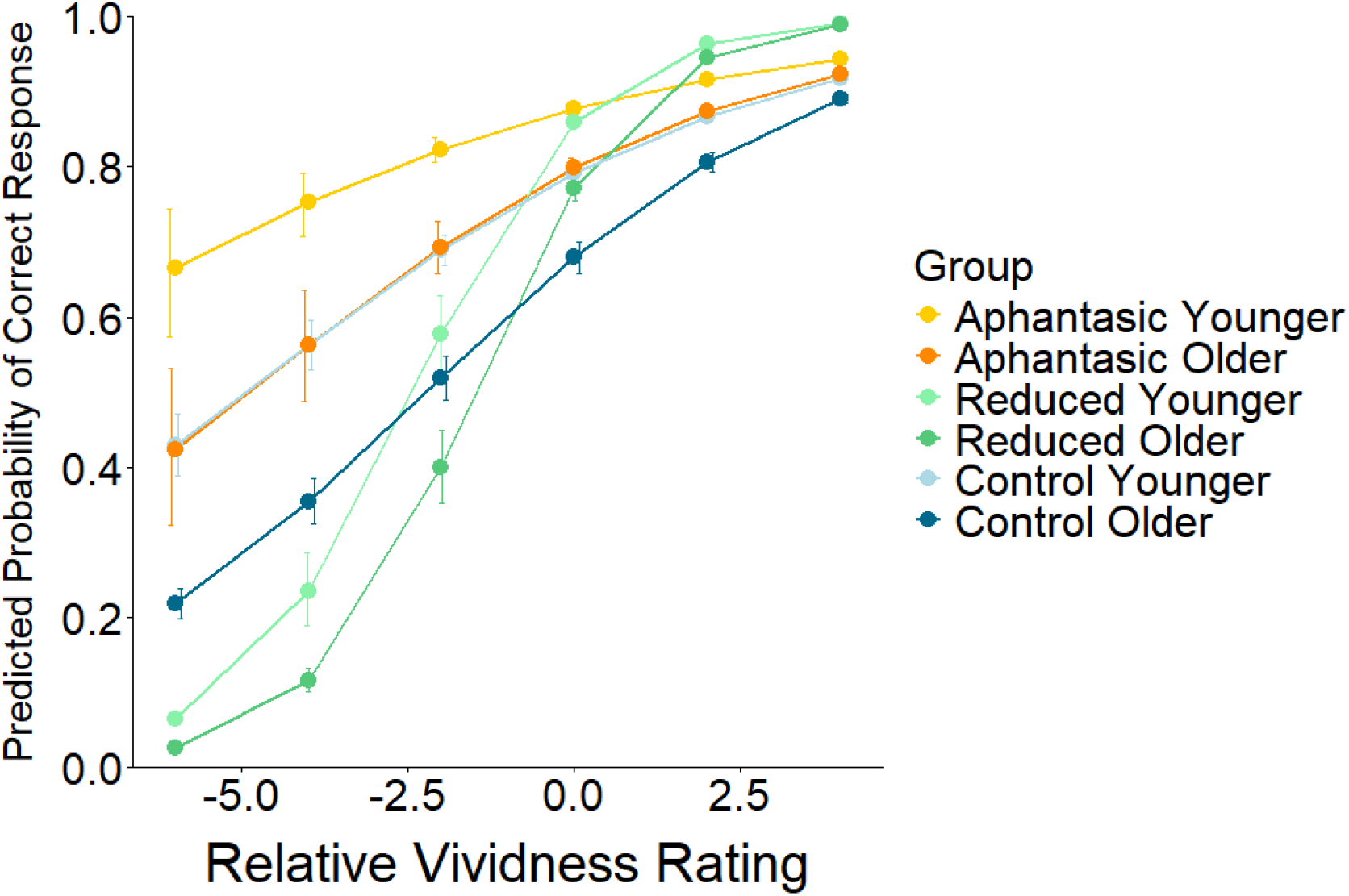
Predicted probabilities of a correct response by Relative Vividness rating. Error bars represent 95% confidence intervals and are offset from means to prevent overlap.

### Confidence

To investigate the effect of imagery group and age category on confidence ratings, a 2×3 two-way ANOVA was conducted. There was a trend towards non-normally distributed residuals following a Shapiro-Wilks test (W = 0.98, *p* = .054), and the data failed a Levene’s test for homogeneity of variance (F(5, 110) = 2.41, *p* = .041), therefore non-parametric ART ANOVA results are also reported to validate parametric findings. Two-way ANOVA analyses revealed a significant main effect of imagery group on confidence ratings (F(2, 110) = 11.40, *p* = <.001), with a non-significant main effect of age category (F(1, 110) = 1.31, *p* = .256), indicating that confidence ratings significantly differed across imagery groups, but not age categories. A significant group x age category interaction was found (F(2, 110) = 5.71, *p* = .002), suggesting the effect of age depended on imagery group. A non-parametric ART ANOVA confirmed findings, with the effects of group (F(2, 110) = 9.52, *p* = <.001), age category (F(1, 110) = 0.89, *p* = .347), and the interaction between group and age category (F(2, 110) = 5.71, *p* = .002), matching reported parametric findings. See Figure 7 for a representation of mean confidence ratings.

**Figure 7.**
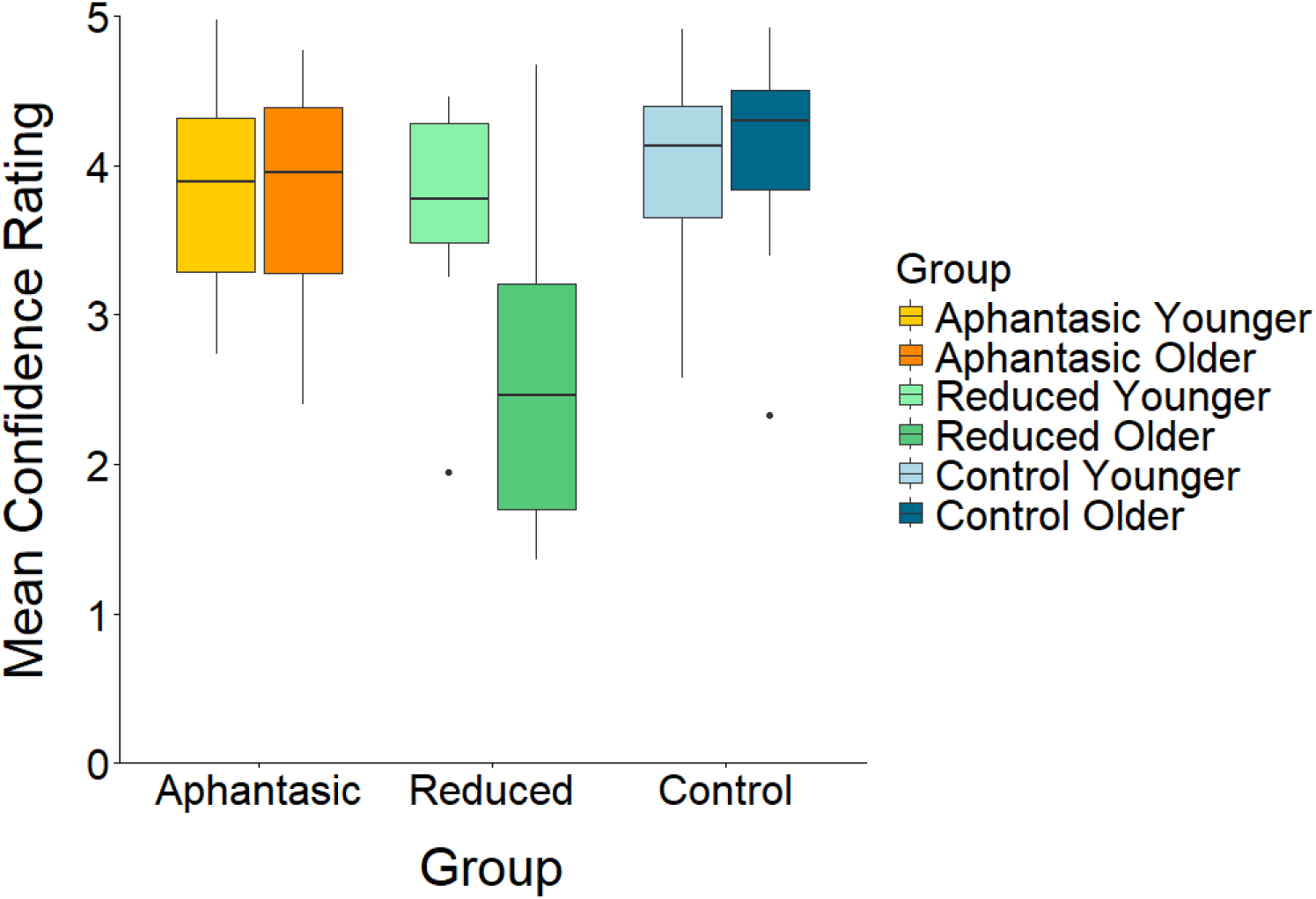
Mean confidence ratings across imagery groups and age categories.

Post-hoc Tukey HSD comparisons for the main Group effect revealed significantly lower confidence ratings between Reduced and Aphantasic (*p* = .008), Reduced and Control (*p* = <.001), but not Aphantasic and Control (*p* = .173) imagery groups, suggesting that, when collapsing across age categories, those with Reduced imagery report significantly lower confidence ratings than those in Aphantasic and Control groups.

To investigate the interaction further, estimated marginal means and pairwise contrasts were conducted within each imagery group. It was found that confidence did not significantly differ between Younger and Older Aphantasic participants (*p* = .835), nor between Younger and Older Control imagery participants (*p* = .504). However, a significant difference between Younger and Older Reduced imagery participants was found (*p* = <.001). Findings were supported by a Wilcoxon rank-sum test for Aphantasic (*p* = .987), Control (*p* = .294), and Reduced (*p* = .020), though the effect in the Reduced group was only a non-significant trend after Bonferroni correction (*p* = .060), therefore, this effect should be interpreted with caution.

### Response Times

To investigate differences in response times when identifying changes in stimuli between imagery groups, a one-way ANOVA was conducted between average response times and imagery group. Data passed a Levene’s test of homogeneity of variance (F(2,113) = 1.69, *p* = 0.189), with response times being normally distributed for the Aphantasic (W = 0.94, *p* = .051) and Reduced (W = 0.94, *p* = .223) imagery groups, but not the Control (W = 0.94, *p* = .004) group. Therefore, non-parametric Kruskal-Wallis and pairwise Wilcoxon rank sum tests are reported to validate parametric results. A one-way ANOVA revealed a significant effect of imagery group on average response time (F(2,113) = 7.51, *p* = <.001). Post-hoc TukeyHSD analyses revealed a significant difference in response times between Control and Aphantasic (*p* = .015) and Control and Reduced (*p* = .003), but a non-significant difference between Aphantasic and Reduced response times (*p* = .578). The main effect of group on response time was confirmed with a Kruskal-Wallis rank sum test (Chi square(2) = 12.71, *p* = .002). Bonferroni-adjusted Wilcoxon tests confirmed faster responses in Controls than in both Aphantasic (*p* = .024) and Reduced (*p* = .007) groups, with no significant difference between the latter two groups (*p* = .960) See Figure 8 for a graphical representation of results.

**Figure 8.**
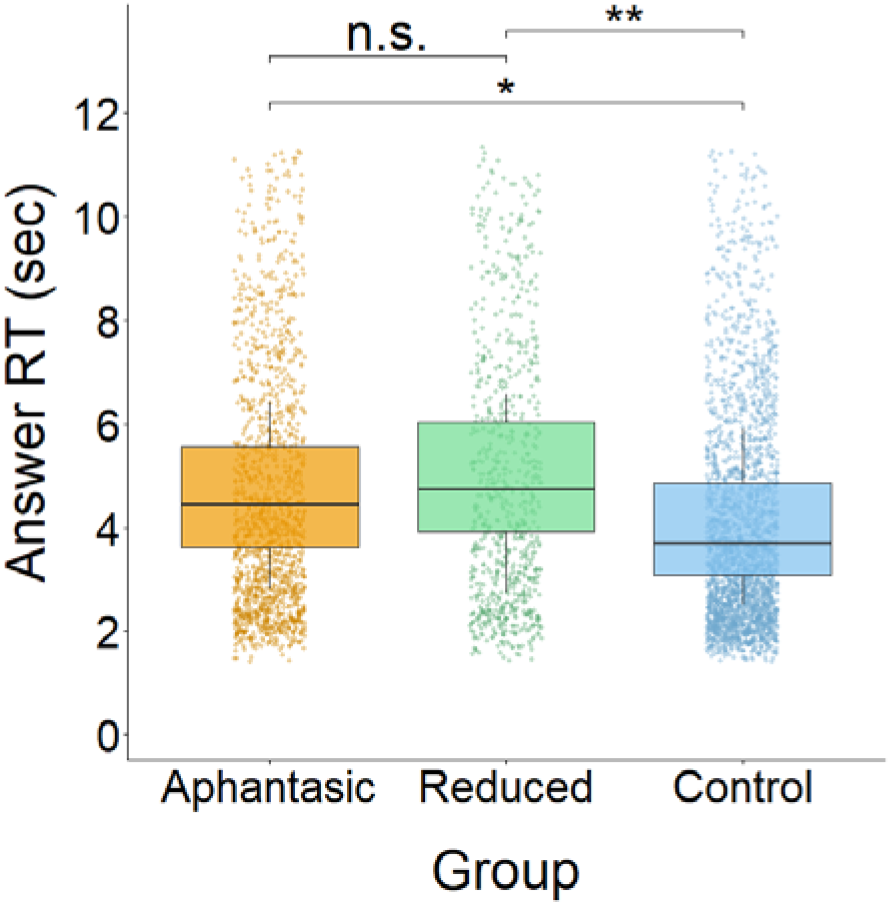
Answer response times by imagery group. Boxplots represent average response times, with jitter points representing raw data points. Significance lines represent pairwise comparisons.

To investigate if the effect of increased response time exists at the state trial-by-trial level, a linear mixed model was fitted predicting log response time (msec) from imagery group, Relative Vividness, and their interaction, with a within-participant random intercept and random slope for Relative Vividness. Note that due to positive skew, log response times were calculated which matched significances as when using raw response times, see Appendix C. See Table 2 for full statistical outputs. Results indicate that at the trial-by-trial level Aphantasic and Reduced participants responded significantly slower than Control participants, with higher Relative Vividness predicting quicker response times across groups. This effect was significantly stronger in the Reduced group compared to Controls, where Aphantasic participants did not significantly differ from Controls.

**Table 2.** Statistical outputs following linear mixed model predicting response times by imagery group, Relative Vividness, and their interaction.

| Variable | Estimate | SE | p |
| --- | --- | --- | --- |
| (Intercept) | 8.16 | 0.03 | <.001*** |
| Group (Aphantasic) | 0.14 | 0.05 | .009** |
| Group (Reduced) | 0.20 | 0.07 | .003** |
| Relative Vividness | -0.70 | 0.01 | <.001*** |
| Relative Vividness * Group (Aphantasic) | -0.01 | 0.03 | .665 |
| Relative Vividness * Group (Reduced) | -0.12 | 0.03 | <.001*** |

Note: Reference group for imagery Group is Control.

To investigate relationships between VVIQ scores and response times when giving answers and vividness ratings, two Spearman’s correlations were carried out as all measures failed Shapiro-Wilks tests for normality. Analyses revealed a significant negative correlation between VVIQ score and average answer response time (r(114) = −0.28, *p* = .003), but a non-significant relationship between VVIQ score and average vividness rating response times (r(114) = −0.10, *p* = .310). Results suggest participants gave their answers quicker the greater their trait vividness was, but this effect failed to be significant when giving vividness ratings, see Figure 9 for a visualisation of this relationship.

**Figure 9.**
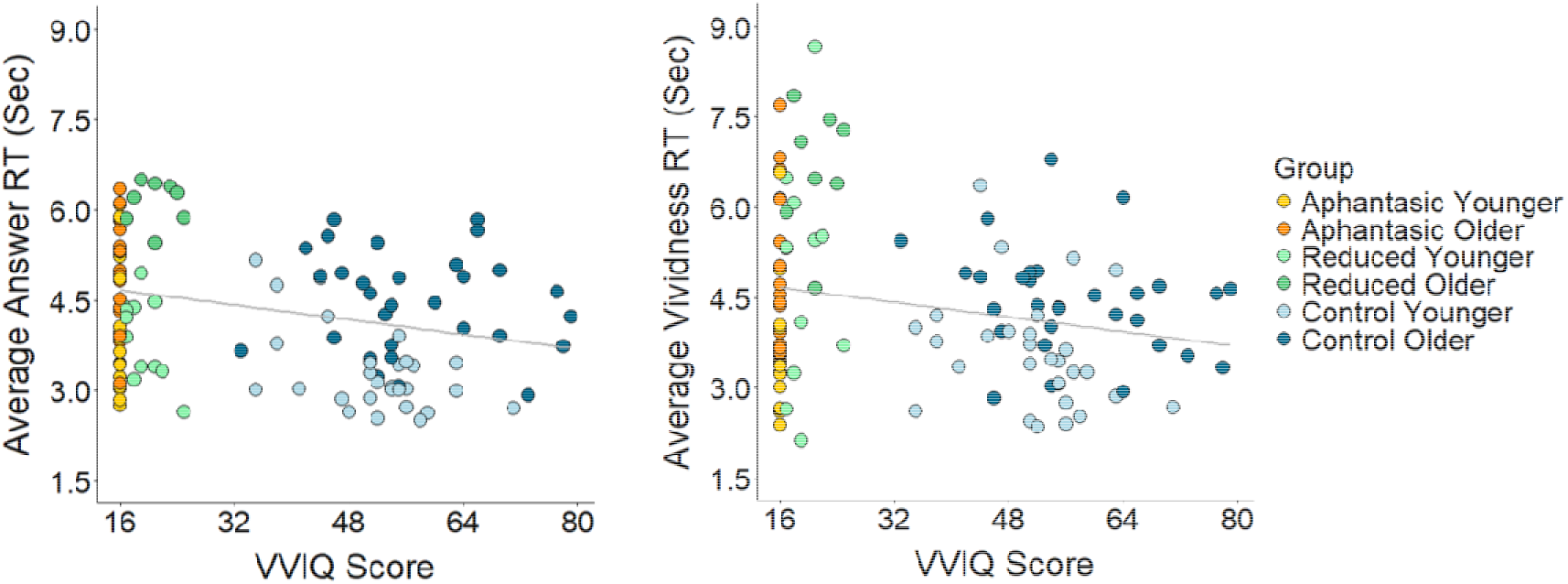
Scatterplot of the relationships between VVIQ scores, and average answer response times (left) and average vividness rating response times (right).

Furthermore, to investigate relationships between trial-by-trial variations in response times and vividness, a linear mixed model was fit to the data with Relative Vividness, imagery group (Control, Aphantasic, Reduced) and their interaction as fixed effects. Random effects included Relative Vividness and participant ID. Response times above the 95^th^ percentile were removed to account for extreme responses. See Table 3 for statistical outputs of the model.

**Table 3.**
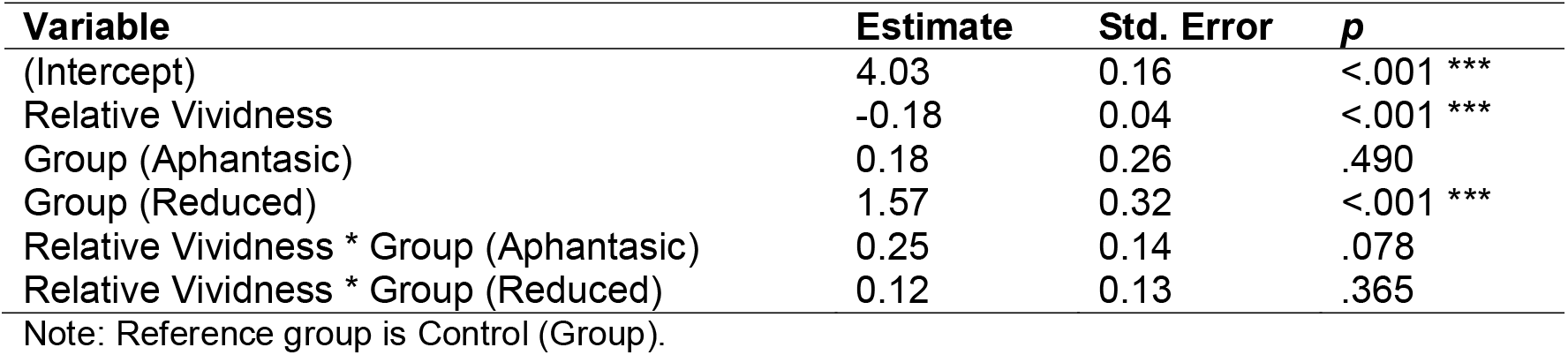
Linear mixed-effects model predicting response time from Relative Vividness ratings and imagery group, and their interaction.

| Variable | Estimate | Std. Error | p |
| --- | --- | --- | --- |
| (Intercept) | 4.03 | 0.16 | <.001 *** |
| Relative Vividness | -0.18 | 0.04 | <.001 *** |
| Group (Aphantasic) | 0.18 | 0.26 | .490 |
| Group (Reduced) | 1.57 | 0.32 | <.001 *** |
| Relative Vividness * Group (Aphantasic) | 0.25 | 0.14 | .078 |
| Relative Vividness * Group (Reduced) | 0.12 | 0.13 | .365 |

Note: Reference group is Control (Group).

Results indicate a significant negative effect of vividness in the Control imagery group, indicating that lower than average vividness ratings were associated with longer response times, and higher than average vividness ratings were associated with quicker response times. Compared to the Control imagery group, those in the Reduced imagery group had significantly longer response times responding on average 1.57 seconds slower than Controls, while those in the Aphantasic imagery group did not significantly differ from Controls. Non-significant interactions between groups and Relative Vividness suggest the relationship between vividness and response time is similar across imagery groups, with the Aphantasic group trending towards significance. See Figure 10 for graphical representations of response times and Relative Vividness for each imagery group.

**Figure 10.**
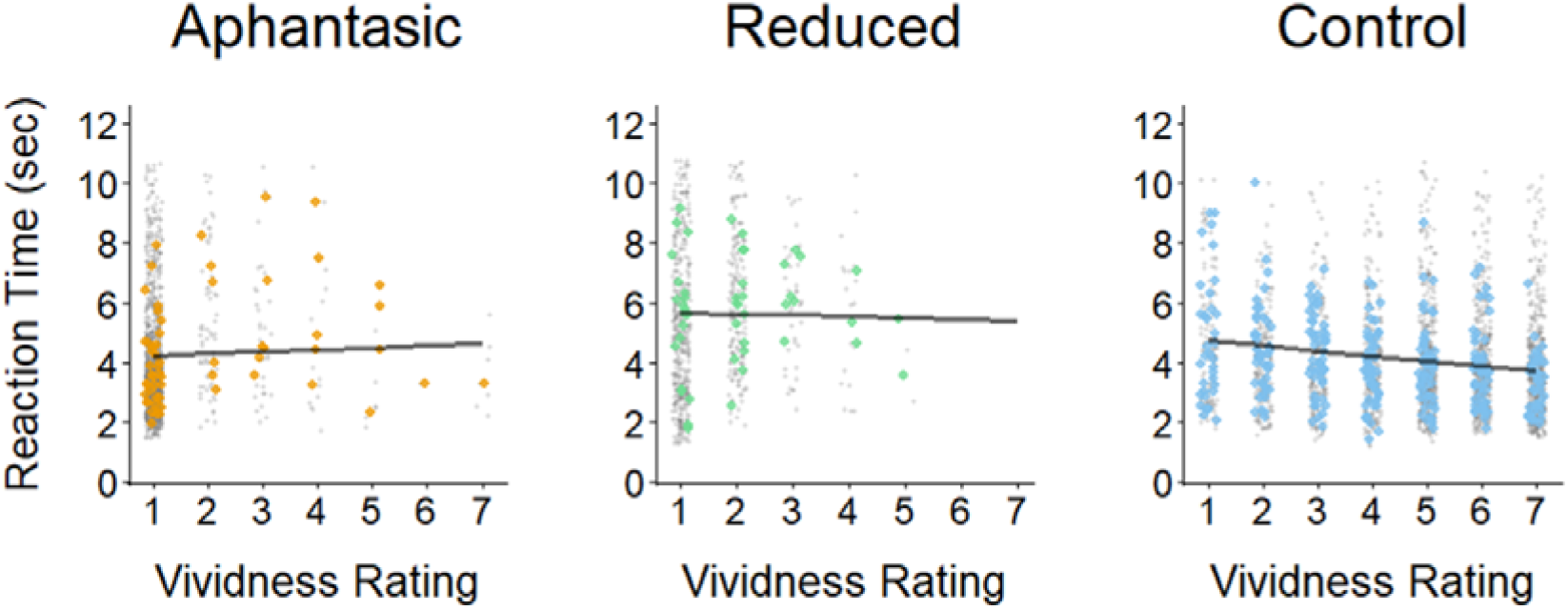
Response time by vividness rating across imagery groups. Coloured dots represent median response times binned to the nearest 0.5 vividness rating. Grey dots represent raw trial-by-trial vividness ratings. Black lines represent group-level model fits following the linear mixed effect model.

### Exit questionnaire data

To gain an understanding of Control participants’ knowledge of aphantasia, a question probing participants’ prior knowledge of aphantasia was included in an exit questionnaire. When asked, 65% (N = 39) of participants in the Control imagery group reported no prior knowledge of aphantasia, with 21.7% (N = 13) reporting some knowledge, and 13.3% (N = 8) reporting prior knowledge of aphantasia. See Figure 11 for a graphical representation of results, and Table 4 for exact response figures.

**Figure 11.**
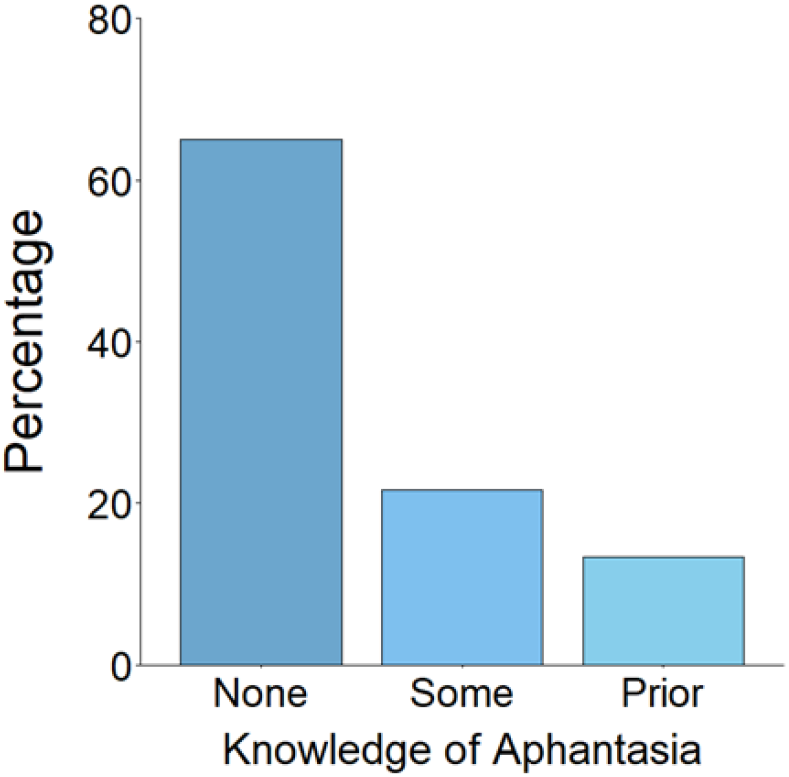
Response percentages of participants with Control imagery ability to the question: “Were you aware of aphantasia prior to this experiment?”

To gain insight into the subjective experience of those with aphantasia and reduced imagery, several questions were included in an exit questionnaire, and word frequency plots of responses were used to ascertain the features that characterised participants’ subjective memory experience. See Appendix A for a full list of questions. Note that Reduced and Aphantasic groups were separated out following data collection, therefore results of questionnaires reflect answers of these two groups combined. The number of respondents to the free text questions were below that necessary for formal qualitative analyses, therefore, only descriptive displays of responses are reported. Common stop words, such as “the” and “and”, and words contained in each respective question, were removed from responses before generating word frequency plots. As can be seen in Figure 12, vividness is often characterised by remembrance and imagery, with the inclusion of emotion. See Figure 12 for graphical representations of responses from participants with self-reported aphantasia, and Table 4 for exact response figures.

**Figure 12.**
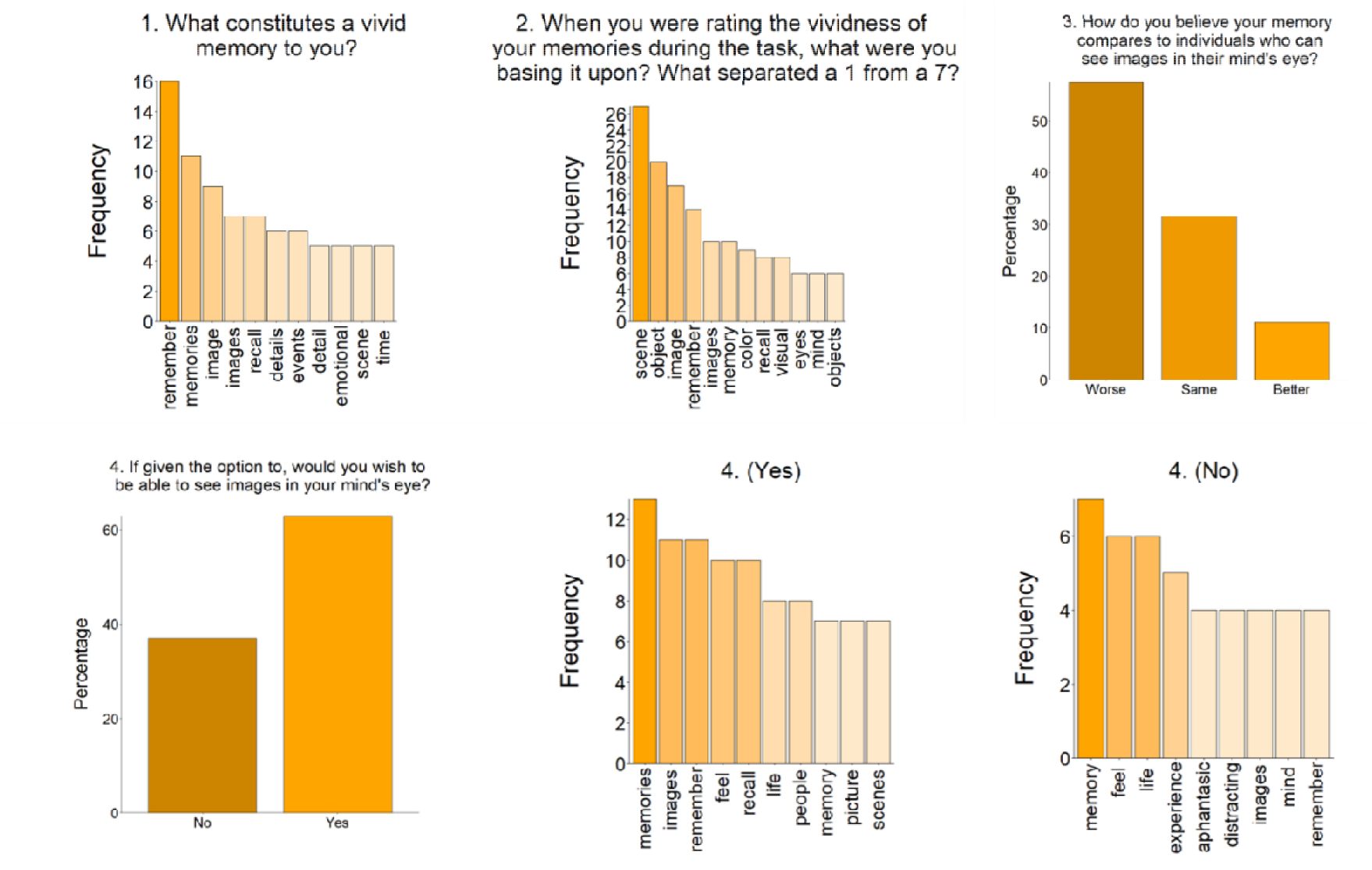
Responses to exit questionnaire questions presented to participants with aphantasia. Table 4.

**Table 4.** Table of response frequencies from given exit questions.

| Question | Response | N | Percentage |
| --- | --- | --- | --- |
| **Control** |  |  |  |
| 1 | Prior knowledge | 8 | 13.3 |
|  | Some knowledge | 13 | 21.7 |
|  | No knowledge | 39 | 65.0 |
| **Aphantasic** |  |  |  |
| 1 |  | 32 |  |
| 2 |  | 32 |  |
| 3 | Better | 6 | 11.1 |
|  | About the same | 17 | 31.5 |
|  | Worse | 31 | 57.4 |
| 4 | Yes | 34 | 63.0 |
|  | No | 20 | 37.0 |
| 4a (Yes – Free Text) |  | 22 |  |
| 4a (No – Free Text) |  | 11 |  |

## Discussion

The present study compared objective memory performance and subjective vividness, confidence, and response times across controls, those with reduced imagery, and aphantasia, in younger and older adults. Trait VVIQ scores and averaged trial-by-trial vividness ratings did not predict objective memory performance. However, trial-by-trial within-subject vividness ratings significantly and positively predicted memory accuracy, an effect that was significantly stronger in the Reduced mental imagery group than in Control and Aphantasic imagery groups. Controls outperformed both Reduced and Aphantasic groups on visual, but not spatial, modality changes. Those with Reduced imagery demonstrated lower confidence than both Aphantasic and Control groups, with a significant difference in confidence between Younger and Older Reduced participants. Both Reduced and Aphantasic groups demonstrated significantly longer response times than Controls when identifying changes, with those in the Reduced group taking significantly longer than both other groups when giving vividness ratings specifically. Younger participants outperformed Older participants overall, though this effect was absent in the Aphantasic group, and the relationship between vividness and accuracy did not differ by age.

### Memory and Vividness

Correlational analyses found no relationship between memory performance and either VVIQ or averaged trial-by-trial vividness ratings, despite a strong correlation between these two between-participant measures. This null relationship has been observed previously (Duckett & Simons, 2026; Knight et al., 2022; Reeder et al., 2024; Siena & Simons, 2024), supporting *H2*, and indicates that considering vividness to be a stable characteristic fails to capture the contribution of vividness during memory retrieval. Following GLMM analyses, which account for trial-by-trial within-subject variations in vividness, a significant positive relationship was found, with higher-than-average vividness on a given trial being associated with a greater probability of being correct, supporting *H1*. Crucially, VVIQ and average vividness ratings were non-significant in the same model. As reported in several studies and meta-analyses, the VVIQ has poor predictive value (Dijkstra & Fleming, 2023; Duckett & Simons, 2026; Z. Liu et al., 2025; Runge et al., 2015, 2017), with the current findings supporting this pattern, and suggesting that trait and averaged measures fail to capture the effect of vividness, which appears to function at the state level. Therefore, the VVIQ may be better suited to categorising individuals by imagery group than to predicting trial-level memory performance.

The significant positive Relative Vividness x Reduced interaction when predicting memory accuracy indicates that trial-by-trial vividness was a stronger predictor of memory accuracy for those with Reduced imagery, than for Control or Aphantasic groups (see Figure 6). This effect was mirrored in response times, where trial-by-trial state vividness was a significantly stronger predictor of response time in the Reduced group than in Controls, where Aphantasics did not significantly differ. One possible interpretation is that imagery may act as a threshold marker for this group, as when a memory is sufficiently strong to generate imagery in someone who infrequently experiences it, imagery itself may signal accuracy, where an absent image signals uncertainty. For controls, imagery is common enough that its presence is less indicative of accuracy, and for aphantasics, imagery is presumed to be absent regardless of accuracy, reducing imagery’s predictive value in both these cases. What constitutes as “vivid” for those with aphantasia, however, is not fully understood, as the Aphantasic group demonstrated variation in vividness ratings (see Figure 10). This raises the possibility that some aphantasic participants used the vividness scale to rate non-visual features of retrieval, such as familiarity, semantic detail, or confidence, rather than visual imagery itself, despite the instructions given. Such an account might explain how 4 self-reported aphantasics exceeded VVIQ scores of 32 in the present study, and the discrepancy between self-reported and VVIQ-defined aphantasia (Beran et al., 2023; Takahashi et al., 2023). What constitutes a “vivid” experience for those with aphantasia remains an open question, requiring further systematic investigation to understand the subjective experience of aphantasia.

### Modality

No group differences emerged for spatial trials, failing to support a spatial advantage for Aphantasics or those with Reduced imagery in longer term memory and failing to support *H3*. While equivalent spatial memory performance has been found between controls and those with aphantasia across a large proportion of the literature (Crowder, 2018; Dawes et al., 2020; Keogh et al., 2021; Pounder et al., 2022; Reeder et al., 2024; Siena & Simons, 2024), the present findings fail to reproduce the spatial advantage for aphantasics reported by others (Bainbridge et al., 2021; L. Kay et al., 2024), suggesting the absence of mental imagery neither impairs nor improves long-term spatial memory. However, Controls identified visual changes significantly more accurately in comparison to Reduced and Aphantasic imagery groups, supporting *H4*. This need not imply a general impairment in those without typical imagery; rather, it suggests that conscious access to visual imagery may provide an additional benefit when identifying visual details. This is consistent with reduced visual detail in aphantasic recall in the literature (Bainbridge et al., 2021; Dawes et al., 2022; Monzel et al., 2022). It also broadly aligns with the blindsight analogy proposed by Michel et al. (2025) where, just as blindsight patients process perceptual visual information without conscious awareness, aphantasic individuals may retain sufficient visual representation without conscious access to mental imagery, but with less accuracy than those with typical imagery. Therefore, access to visual mental imagery appears to be beneficial for long-term visual memory, with no evidence to suggest a lack of imagery improves or hinders spatial memory.

### Response Times

Controls responded significantly faster than both other groups when identifying changes, supporting *H5*, with no significant difference between Reduced and Aphantasic groups’ response times. Furthermore, higher trial-by-trial Relative Vividness was a significant predictor of shorter response times, with this effect being stronger in the Reduced group when compared to Controls, with no difference in the strength of this effect between Aphantasics and Controls. Together with a significant negative correlation between VVIQ and average response times, these results extend previous findings of slower responding in aphantasia from mental rotation (L. Kay et al., 2024; Pounder et al., 2022), imagination-perception (J. Liu & Bartolomeo, 2023), and visual working memory tasks (Keogh et al., 2021) to long-term memory. This suggests that the compensatory non-visual strategies used during short-term memory tasks may also be utilised in long-term memory, with similar response time costs. The absence of a difference between Reduced and Aphantasic groups is notable, as Pounder et al. (2022) found response time differences only after excluding reduced imagery participants from total aphantasics, suggesting that the reduced imagery participants in their sample skewed results with quicker, Control-like, response times. The present findings may reflect different cognitive demands during long-term retrieval, with the uncertainty in weak imagery potentially leading to increased hesitation or effort, resulting in increased response times. This interpretation warrants further investigation with increased sample sizes to determine the exact nature of response times between imagery groups, and across different memory contexts.

Regarding vividness ratings specifically, higher ratings were given more quickly across all groups supporting *H7* and extending Barnett et al. (2026) to aphantasic and reduced imagery groups. Notably, the Reduced group took significantly longer than both Controls and Aphantasics when rating vividness, while Aphantasics and Controls did not differ from one another. The presumed absence of mental imagery in aphantasics likely resulted in quick responses of low vividness in the present sample, and longer responses to determine a “vivid” response. Those with reduced imagery however, take longer to assess their own sense of vividness on each trial, perhaps in part due to greater metacognitive effort being required to determine whether an image is present, and if so, how clear it may be. Therefore, increased vividness response times, alongside task response times, appear to be behavioural markers of Reduced imagery, possibly due to metacognitive uncertainty.

### Confidence

Confidence did not differ significantly between Aphantasic and Control groups, failing to support *H6*, and prior findings in the literature (Monzel, Dance, et al., 2023; Wittmann & Şatırer, 2022). Instead, those in the Reduced imagery group reported significantly lower confidence than both other groups. One explanation is that prior studies have collapsed low VVIQ scorers into a single aphantasic group, leading the confidence gap to be attributed to aphantasia, whereas it may instead be driven primarily by those with weak access to mental imagery, obscuring effects as suggested by Phillips (2026). As the confidence gap may be attributable to uncertainty in one’s mental imagery, the Reduced group, with weak and inconsistent access to mental imagery, may have lower confidence in their answers as a result, where those with Aphantasia appear to have a less pronounced reduction in confidence. Present findings extend the absence of a confidence gap between aphantasics and controls during working memory (Reeder et al., 2024) to long-term episodic memory. This may be due to the nature of the task in the present study, as participants were presented with the scene image again during the test phase, with or without a change, rather than requiring participants to recreate and rely solely on the image within their mind’s eye. In this context, confidence may not have varied sufficiently to reveal group differences. Further research is required, with larger sample sizes and more varied and complex episodic memory tasks to investigate any potential confidence gaps between imagery groups within long-term memory.

A significant Group x Age interaction indicated that Older Reduced imagery participants were less confident than the Younger Reduced. One possible interpretation is that older adults with weak but present imagery experience a compounding of two sources of metacognitive uncertainty; age-related memory decline as described in the broader aging literature (Folville et al., 2021) and the presence of weak and perhaps unreliable mental imagery. Combining these two factors may contribute to greater uncertainty in one’s memory, lowering subjective confidence. Alternatively, the Older Reduced group was the smallest sample, with N = 8, therefore it may be that the sample contained, by chance, several unconfident individuals giving rise to this effect. Therefore, larger sample sizes of those with reduced imagery should be obtained in future studies to confirm or refute findings.

### Reduced imagery as a distinct group

Across the present analyses, those with Reduced, but not totally absent imagery (VVIQ = 17-32), emerge as a distinct group from both Aphantasic (VVIQ = 16) and Control (VVIQ > 32) groups in three ways: a significantly stronger effect of trial-by-trial vividness on accuracy (see Figure 6) and response times (see Table 2), significantly longer vividness rating times, and significantly lower confidence. Taken together, these findings suggest that reduced imagery may be characterised by a more effortful and uncertain imagery experience distinct from typical imagers’ fluent conscious access to imagery, and those with aphantasia using alternative non-visual strategies. Reduced imagery has been referred to as “hypophantasia” (Reeder et al., 2024), with others supporting the notion that combining aphantasia and reduced imagery together obscures effects (Phillips, 2026). For example, group differences have been found during reasoning (Le Clézio et al., 2026), implicit priming (Purkart et al., 2025), and mental health outcomes (Kvamme et al., 2026) as those with reduced imagery “may differ fundamentally” from aphantasics (Purkart et al., 2025). The profile of increased uncertainty may be relevant to recent findings linking reduced imagery to poorer mental health outcomes (Kvamme et al. (2026), although the present study was not designed to test that relationship directly. Critically, the present findings suggest that collapsing all low scorers on the VVIQ into a single aphantasic group may obscure specific effects unique to those with reduced imagery, which may explain certain inconsistencies in the literature. Future research should therefore consider those with reduced, but not totally absent mental imagery, as a group distinct from both total aphantasics and controls.

### Age-related effects

Younger participants demonstrated greater memory performance than Older participants overall, supporting *H8*, with significant within-group differences in Control and Reduced imagery groups, but a non-significant difference between Younger and Older Aphantasics (See Figure 5). Although the Group x Age interaction was non-significant, and the subsample of Aphantasics was relatively small, the findings raise the possibility that the cognitive strategies employed by those with aphantasia are less susceptible to age-related decline than imagery-based strategies. If individuals with aphantasia rely more on semantic, verbal, or other non-visual strategies, which may be relatively preserved in healthy aging (Hedden & Gabrieli, 2004), this could attenuate the age-related decline in memory observed in individuals with typical or reduced imagery. However, this interpretation remains speculative but represents a possibility that future research should seek to investigate with larger samples.

The non-significant GLMM interaction between Age Category and Relative Vividness indicates vividness contributed similarly to accuracy for Younger and Older participants, failing to support *H9*. This finding is consistent with Mojescik et al. (2024), who found that both age groups anchored their vividness ratings to memory gist in a comparable way, suggesting that state-level vividness-memory relationships may be relatively stable across the lifespan. Subjective vividness, at the state level, appears to reflect retrieval success in parallel to objective memory performance, indicating that the cognitive mechanisms facilitating this relationship are closely related.

### Subjective experience of reduced and absent imagery

Word frequency analyses of exit questionnaire responses provided insight into what individuals with aphantasia and reduced imagery consider a “vivid” memory. While many responses reflected the conventional view of vividness (a clear, distinct image akin to real life), others defined it through emotional connection, linguistic representations, and a sense of “knowing”, suggesting that, for some participants, vividness ratings may have reflected non-visual aspects of retrieval. This further highlights the heterogeneity of aphantasia, as participants appeared to use the term “vividness” in different ways, which may explain why vividness had less predictive value for our total Aphantasic sample in comparison to the Reduced sample.

Despite no overall objective memory deficit between groups in the present study, when asked, 57.4% of aphantasic sample believed their memory to be worse than that of typical imagers. This is consistent with the confidence gap noted in the literature (Monzel, Dance, et al., 2023), which persists independently of actual performance. The present findings do not support this belief in the context of the current memory task, yet it is held by the majority of the sample, highlighting a discrepancy between the subjective experience and objective performance of those who have reduced or absent imagery. Furthermore, when asked whether they would want visual imagery given the option, 63% said yes, frequently citing improvements to objective memory performance and improved subjective qualities during recall. Of the 37% who would not choose to see images, respondents cited concerns over changes in cognition and ways of thinking, distractions, and unwanted changes to their sense of self (See Digard et al. (2026)). Assumptions that imagery is a desirable quality of subjective experience holds for the majority of our sample, but a substantial proportion of the same sample did not wish to obtain mental imagery. These findings highlight a lack of informed understanding about mental imagery and its contribution to memory performance among aphantasics, demonstrating the need for both increased dissemination of findings, and further research.

### General knowledge of aphantasia

Among control participants, 65% reported no prior knowledge of aphantasia. This is an important finding, as it suggests that the general population may be largely unaware of individual differences in subjective experience, which may also influence prevalence estimates. This lack of awareness may reduce the proportion of people who self-identify as aphantasic and therefore complicate estimates of self-reported prevalence. Prevalence of aphantasia varies in the literature, from 1% (Dance et al., 2022; Zeman, 2024) for total aphantasics, to as high as a self-reported prevalence of 8.9%-12.1% (Beran et al., 2023; Takahashi et al., 2023). As reported by Zeman et al. (2020) over 60% of individuals with aphantasia discover aphantasia after the age of 20 following discussions with friends and family, media exposure, and/or visualisation practices. Together, these results suggest that aphantasia and mental imagery are rarely discussed within the general population, and awareness of aphantasia emerges relatively late in life despite representing an extreme variation in subjective experience and a key aspect of one’s sense of self (Digard et al., 2026). Therefore, disseminating research findings is important to increase knowledge of mental imagery in the general population, and has the potential to improve prevalence estimates of those with aphantasia.

## Conclusion

The present study supports a clear distinction between typical, reduced, and absent mental imagery. Individuals with reduced imagery demonstrate a profile characterised by a greater contribution of vividness to episodic memory accuracy, longer response times when rating vividness, and lower confidence in their answers. Additionally, between-participant measures of vividness fail to predict memory performance across groups, while trial-by-trial state measures of vividness are significant positive predictors of accuracy, indicating the effect of vividness on memory operates at the state level. Controls outperformed both other groups on visual changes specifically, with equivalent spatial performance across groups, suggesting conscious access to imagery specifically benefits visual long-term memory. Aphantasic and Reduced groups took significantly longer to give their answers than Controls. No confidence differences were found between Control and Aphantasic groups, contrary to prior literature, but reduced confidence was instead found in the Reduced group, indicating metacognitive uncertainty in inconsistent imagery reduces confidence. Younger participants outperformed Older participants in Control and Reduced groups, with no significant difference in the Aphantasic group. Finally, aphantasia appears to be a heterogeneous subjective experience, characterised by marked reductions in perceived memory ability, while the general population appears to be largely unaware of individual differences in mental imagery. Taken together, the findings highlight the importance of measuring vividness at the trial-specific level, distinguishing reduced imagery from total aphantasia, and further investigating the subjective experience of reduced and absent mental imagery.

## Author contributions

W.D. contributed to conceptualisation, methodology, investigation, data curation, formal analysis, visualisation, and writing the original draft. J.S.S. contributed to conceptualisation, methodology, supervision, funding acquisition, and writing - review and editing. Both authors approved the final manuscript.

## Acknowledgements

This research was supported by the UK Research and Innovation (UKRI) Economic and Social Research Council and by a UKRI Cross Research Council Responsive Mode award [MR/Z505614/1]. We would like to thank Tom Ebeyer from *Aphantasia Network* for his assistance during participant recruitment. For the purpose of open access, the author has applied a Creative Commons Attribution (CC BY) licence to any Author Accepted Manuscript version arising from this submission.

## Declaration of competing interest

The authors declare no competing interests.

## Data availability

Anonymised data are available on OSF (https://osf.io/93g8j/overview?view_only=3d7147be777c402fbc61a639c34ed592)

## Appendix

## Appendix A. Exit questionnaire questions for the Control, and Aphantasic and Reduced groups.

Control questions:

1. Were you aware of aphantasia prior to this experiment? (Multiple choice)

1. Yes, I was aware of what aphantasia was and what it meant
2. I had a vague knowledge of aphantasia, but wasn’t sure exactly what it was
3. I had not heard of aphantasia prior to this experiment

Aphantasic and Reduced questions:

1. What constitutes a vivid memory to you? (Free text)
2. When you were rating the vividness of your memories during the task, what were you basing it upon? What separated a 1 from a 7? (Free text)
3. How do you believe your memory compares to individuals who can see images in their mind’s eye? (Multiple choice)

1. Better
2. About the same
3. Worse
4. If given the option to, would you wish to be able to see images in your mind’s eye? (Multiple choice)

1. Yes, I would want to be able to ‘see’ images in my mind’s eye
2. No, I would prefer to be aphantasic

4a Why, or why not? (Free text)

## Appendix B. Frequency of each Relative Vividness rating for each imagery group, rounded to the nearest 0.5

|  | -5.5 | -5 | -4.5 | -4 | -3.5 | -3 | -2.5 | -2 | -1.5 | -1 | -0.5 | 0 | 0.5 | 1 | 1.5 | 2 | 2.5 | 3 | 3.5 |
| --- | --- | --- | --- | --- | --- | --- | --- | --- | --- | --- | --- | --- | --- | --- | --- | --- | --- | --- | --- |
| **Aphantasic** | 0 | 0 | 0 | 0 | 0 | 3 | 0 | 7 | 15 | 40 | 44 | 1139 | 35 | 18 | 6 | 4 | 5 | 8 | 1 |
| **Control** | 1 | 5 | 10 | 15 | 18 | 51 | 79 | 114 | 153 | 161 | 264 | 322 | 371 | 354 | 208 | 123 | 61 | 27 | 13 |
| **Reduced** | 0 | 0 | 0 | 0 | 0 | 0 | 0 | 4 | 4 | 22 | 177 | 283 | 173 | 43 | 21 | 5 | 0 | 1 | 0 |

## Appendix C. Results of the linear mixed effects model predicting trial-by-trial response times (msec) from imagery group and Relative Vividness, using raw Relative Vividness ratings.

| **Variable** | **Estimate** | **SE** | ***p*** |
| --- | --- | --- | --- |
| (Intercept) | 3945.56 | 137.59 | <.001*** |
| Group (Aphantasic) | 631.97 | 225.10 | .006** |
| Group (Reduced) | 930.61 | 275.82 | .001** |
| Relative Vividness | -263.12 | 39.31 | <.001*** |
| Relative Vividness * Group (Aphantasic) | -118.42 | 148.20 | .427 |
| Relative Vividness * Group (Reduced) | -561.01 | 144.50 | <.001*** |

## Notes

https://osf.io/93g8j/overview?view_only=3d7147be777c402fbc61a639c34ed592

### Competing Interest Statement

The authors have declared no competing interest.

